# Computational structural prediction and chemical inhibition of the human mitochondrial pyruvate carrier protein heterodimer complex

**DOI:** 10.1101/2024.05.16.594520

**Authors:** Christy M. Hadfield, John K. Walker, Chris Arnatt, Kyle S. McCommis

**Affiliations:** Edward A. Doisy Department of Biochemistry & Molecular Biology, Saint Louis University School of Medicine; Department of Pharmacology & Physiology, Saint Louis University School of Medicine; Department of Chemistry, Saint Louis University

**Keywords:** mitochondrial pyruvate carrier, pyruvate transporter, mitochondria, thiazolidinedione, UK-5099

## Abstract

The mitochondrial pyruvate carrier (MPC) plays a role in numerous diseases including neurodegeneration, metabolically dependent cancers, and the development of insulin resistance. Several previous studies in genetic mouse models or with existing inhibitors suggest that inhibition of the MPC could be used as a viable therapeutic strategy in these diseases. However, the MPC’s structure is unknown, making it difficult to screen for and develop therapeutically viable inhibitors. Currently known MPC inhibitors would make for poor drugs due to their poor pharmacokinetic properties, or in the case of the thiazolidinediones (TZDs), off-target specificity for peroxisome-proliferator activated receptor gamma (PPARγ) leads to unwanted side effects. In this study, we develop several structural models for the MPC heterodimer complex and investigate the chemical interactions required for the binding of these known inhibitors to MPC and PPARγ. Based on these models, the MPC most likely takes on outward-facing (OF) and inward-facing (IF) conformations during pyruvate transport, and inhibitors likely plug the carrier to inhibit pyruvate transport. Although some chemical interactions are similar between MPC and PPARγ binding, there is likely enough difference to reduce PPARγ specificity for future development of novel, more specific MPC inhibitors.

## INTRODUCTION

The human mitochondrial pyruvate carrier (MPC) is responsible for transporting pyruvate across the impermeable inner mitochondrial membrane. This critical checkpoint links cytosolic glycolysis and lactate metabolism to the tricarboxylic acid (TCA) cycle, thereby providing pyruvate carbons for both oxidative phosphorylation as well as anaplerotic and biosynthetic reactions that initially occur within the mitochondrial matrix ^[1]^ **(Figure 1)**. Due to this nearly universal importance, the MPC plays a critical role in many diseases including cancers like colon, brain, breast, and liver cancer ^[2]^, neurodegeneration in Alzheimer’s and Parkinson’s ^[3]^, and the development of insulin resistance ^[4]^. This insulin resistance can lead to many other diseases, including diabetes, metabolic dysfunction-associated steatotic liver disease (MASLD), and cardiovascular disease associated with obesity.

**Figure 1:**
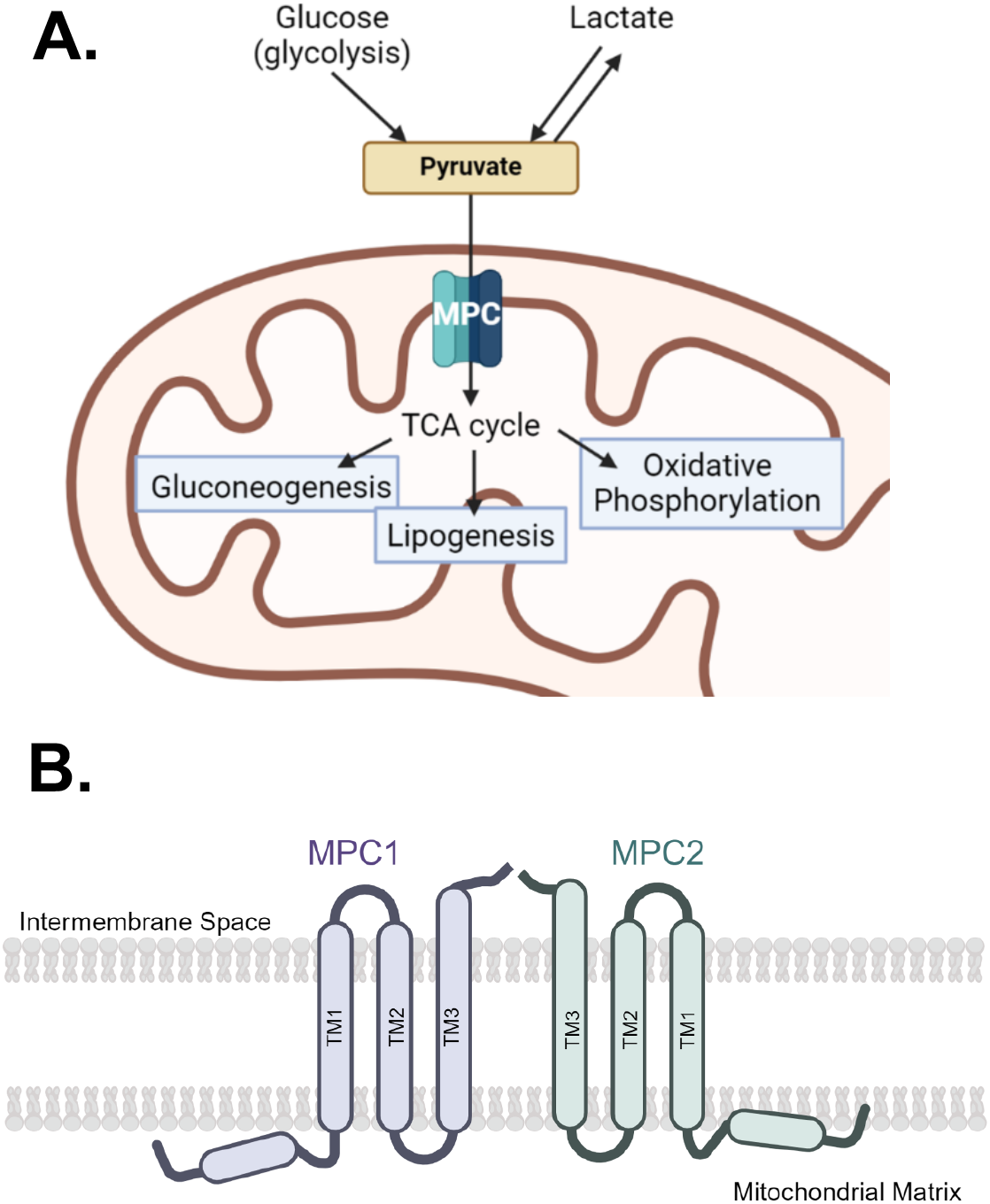
MPC’s role in metabolic pathways. **A**. The mitochondrial pyruvate carrier (MPC) is an inner mitochondrial membrane protein that transports pyruvate into the mitochondrial matrix. From there, pyruvate carbons can enter the TCA cycle and provide substrates for oxidative phosphorylation or be used in anaplerotic/biosynthetic reactions. **B**. The human MPC is a heterodimer comprised of MPC1 and MPC2 monomers. Although much is unknown about the structure and orientation of MPC within the inner mitochondrial membrane, it is suspected that each monomer is comprised of three transmembrane domains and an amphipathic N-terminal helix. Figure created with BioRender.com.

Reports of humans with MPC mutations show very severe phenotypes, supporting the functional importance of this protein ^[5], [6], [7], [8]^. Global knockout of MPC in mice is lethal in the embryonic stages ^[9], [10], [11], [12]^, but liver-specific Mpc deletion in mice has shown beneficial metabolic effects ^[13], [14], [15], [16]^, leading to an interest in developing an MPC inhibitor as a possible therapeutic ^[3]^. However, a significant hurdle in the discovery and development of MPC-specific inhibitors is that the protein structure of the MPC complex is unknown. Because the MPC is a small membrane protein, experimental structural determination using common methods such as X-ray crystallography and Cryo-EM is difficult.

Although the structure of the MPC complex is unknown, some key features of the complex have been identified. Specifically, the functional unit of the human MPC is a heterodimer made up of MPC1 and MPC2 proteins. It is suspected that each monomer contains three transmembrane domains and an N-terminal amphipathic helix ^[4]^ **(Figure 1B)**. What is not known, however, is the conformation of the monomers and their complex, functionally important residues during pyruvate transport, and the structural dynamics of the protein during transport. Additionally, while MPC1 is ~11kDa and MPC2 ~14kDa, comprising a ~25kDa heterodimer, the finding of ~150kDa complex in native gel electrophoresis ^[17]^ suggests oligomerization may occur. It has also been widely accepted that although they do not share much sequence identity, the MPC heterodimer is structurally similar to the bacterial SemiSWEET sugar transporter, allowing for the creation of homology models to hypothesize the structure of the MPC heterodimer ^[18]^.

Several classes of small molecules can bind and inhibit the MPC, such as UK-5099, an α-cyano-cinnamate derivative ^[19]^, the anti-cancer agent 7ACC2, originally thought to inhibit plasma membrane monocarboxylate transporters ^[20]^, and the insulin-sensitizing thiazolidinedione (TZD) compounds **(Table 1)**. TZDs are also agonists for the nuclear receptor peroxisome-proliferator-activated receptor gamma (PPARγ) which regulates the expression of genes associated with adipogenesis and fat storage, primarily in adipose tissue.

**Table 1:**
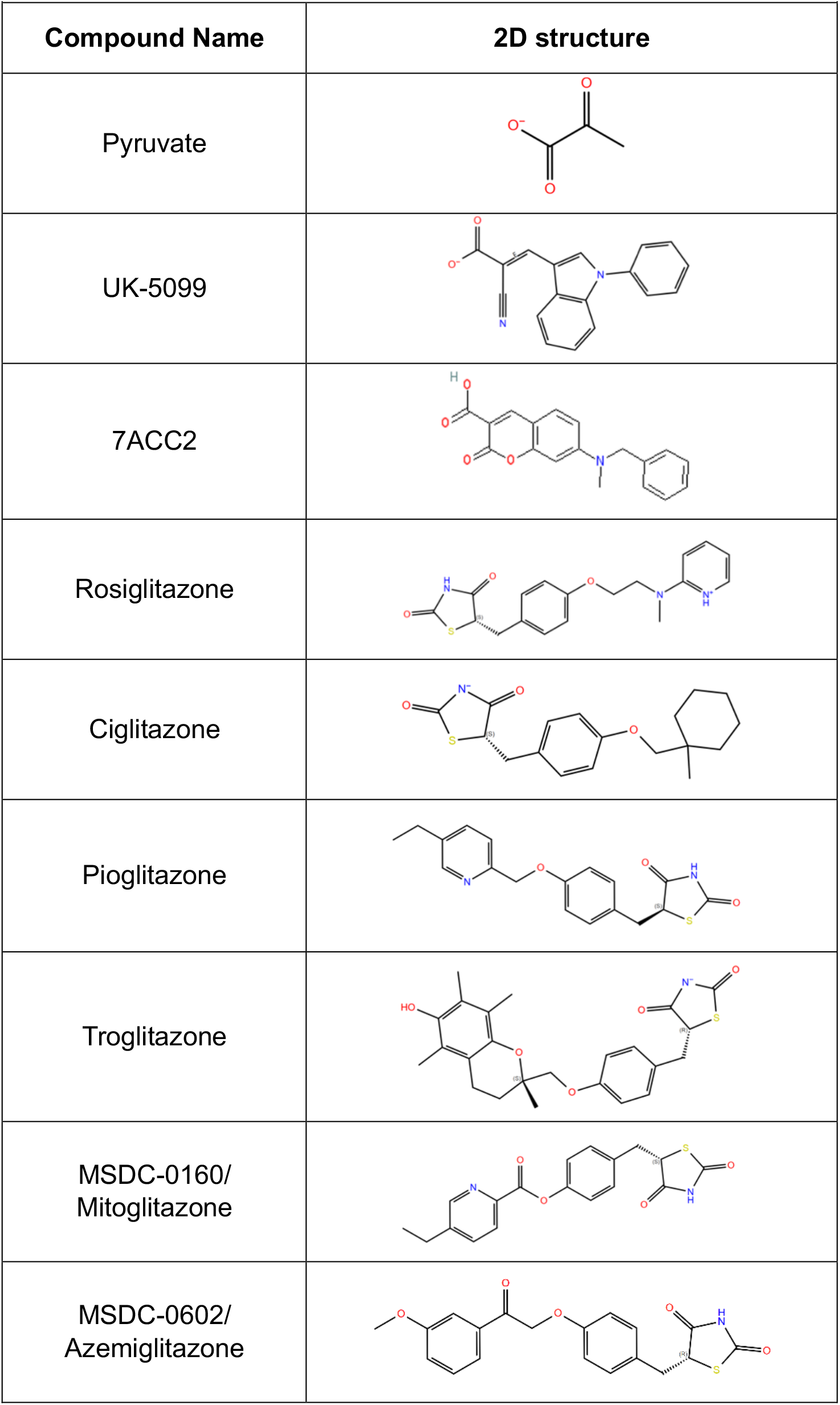
Known ligands and inhibitors of MPC.

Interestingly, different TZD molecules display diverse potencies for PPARγ binding and activation ^[21], [22]^, and pioglitazone, which has superior clinical benefit, displays significantly less PPARγ agonism compared to rosiglitazone or troglitazone ^[1]^. Therefore, it is suspected that the beneficial metabolic effects of TZDs may arise instead from MPC inhibition. MSDC-0160 and MSDC-0602 were developed with this in mind, and display significant antidiabetic benefits despite little-to-no PPARγ activation ^[22], [23], [24], [25], [26], [27]^. Along these lines, PXL065, which is a stabilized R-enantiomer of pioglitazone shown to inhibit the MPC but not bind PPARγ, is also in development for MASLD ^[28], [29]^. MPC inhibitors without an impact on PPARγ would likely make for better insulin-sensitizing compounds while avoiding unwanted side effects.

The alternating access mechanism of transport of the MPC, as proposed for solute transporters by Jardetzky in the 1960s ^[30]^, likely involves a series of protein conformational changes allowing access of the ligand binding site to the mitochondrial intermembrane and matrix spaces. A third occluded (O) state could also exist, possibly upon substrate or inhibitor binding. While it is not currently known if MPC takes on these different conformations, the bacterial SemiSWEET sugar transporter is known to take on both the OF, IF, and an occluded (O) conformation ^[31]^. There have also previously been Bioluminescent Resonance Energy Transfer (BRET) assays developed for the MPC suggesting structural movement within MPC ^[32], [18], [33]^. In these assays, a luciferase photon donor is fused to the C-terminus of MPC2 and a fluorescent protein photon acceptor is C-terminally fused to MPC1. Expressing these constructs in cells and then treating with either pyruvate or inhibitor compounds causes an increase in BRET, indicating the C-terminal photon donor and acceptor move to closer proximity. This suggests a conformational change during pyruvate transport or inhibition. Using this assay, several amino acid residues were identified that when mutated, decreased the ability of pyruvate or inhibitors to increase BRET signal ^[18]^. Those important amino acid residues are MPC1 Phe66, MPC2 Lys49, and MPC2 Asn100. These results suggest that these residues are critical for either binding pyruvate or inhibitors, or for coupling pyruvate/inhibitor binding to the gating and conformational changes.

With the lack of experimentally determined structural information for the MPC, in this current study, we develop a homology model of the human MPC heterodimer modeled off the bacterial SemiSWEET transporter crystal structure in the outward-facing conformation. Additionally, we generated an AlphaFold predicted heterodimer structures to investigate the binding of currently known inhibitors and the different conformational states.

## EXPERIMENTAL DETAILS

### MPC structural model generation

Generation of the structural models of the MPC was done via two different methods. First, we performed homology modeling, which is similar to methods previously used for the MPC ^[18]^. To do this, first, AlphaFold ^[34]^ was used to predict the structure of the hMPC1 and hMPC2 monomers independently. Then TM-Align ^[35]^ was used to map each monomer independently to a protein found with suspected homology, which in this case is the bacterial SemiSWEET sugar transporter in the outward-facing (PBD: 4X5N) or occluded conformation (PDB: 4QNC). The SemiSWEET transporters are homodimers, so MPC1 was mapped to one of the monomer positions, and MPC2 was mapped to the other position. Then, the coordinates provided by TM-Align were used to align the MPC monomers into a heterodimer state in PyMOL ^[36]^.

For the second method, heterodimer models were generated using AlphaFold2 multimer v3 ^[37], [38]^. This program fully predicts the structures of the monomers and how they fit together as a heterodimer. Models were first generated using two different settings: either a template database was not used (group 1), or the PDB structure database was used as a template (group 2). Additionally, these generated models were either relaxed to the lowest energy state using Amber ^[39]^, or unrelaxed.

Molecular dynamics (Desmond, Schrödinger) were used to predict the heterodimer positioning in the membrane ^[40]^. For this system, POPC (300k) lipids and TIP3 water modeling was used, totaling 60,753 atoms building an orthorhombic box. The simulation time was 100ns with 10ps recording intervals, using the NVT ensemble class and a temperature of 310 K. The membrane was relaxed before the simulation.

Transmembrane regions of the monomers were also predicted using the TmAlphaFold Transmembrane Protein Structure Database ^[41]^. Binding pockets and tunnels were visualized and obtained using MOLE*online* ^[42]^, which was also used to assign conformation states to each model.

### Binding site generation in structural models

The Schrödinger Suite software was used for all further modeling and exploration of the models. All protein models and structures were modified/prepped using Protein Prep ^[43]^. The termini were capped, and H-bonds were optimized. The structures were minimized to their lowest energy conformation using the OPLS4 force field via Prime ^[44], [45]^. For our structural models of the MPC, SiteMap was used to generate potential ligand binding pockets for docking ^[46], [47]^. For the PPARγ crystal structure containing a bound rosiglitazone ligand (PBD: 7AWC) ^[48]^, the binding pocket was built around where the ligand was already bound. Generation of the receptor-binding grids was done using Glide ^[49], [50], [51]^. Only the default settings were used, unless otherwise specified.

### Ligand docking

All ligands being investigated, including endogenous ligands, inhibitors, and small molecules in library databases were prepped using LigPrep ^[52]^. Only the default settings were used, unless otherwise specified. Once binding sites were determined, ligands in small quantities were docked directly using Glide ^[49], [50], [51]^. Only the default settings were used, unless otherwise specified. The quality of docking was analyzed using Glide Scores measuring free energy of binding. For sets of known ligands and inhibitors, extra precision (XP) settings were used.

### Figure generation

Structures generated by the above methods were polished in PyMOL ^[36]^ to create graphical figures. The graphical representation of amino acids involved in binding was created using GraphPad Prism version 10.0.0^[53].^

## EXPERIMENTAL RESULTS & DISCUSSION

### MPC structural models

The structure of human MPC has yet to be discovered using traditional protein structural determination methods such as X-ray crystallography or Cyro-EM, which is likely due to the difficulty of working with small and dynamic heteromeric membrane proteins. To perform computational docking of compounds, a structural model of the MPC heterodimer was needed, which was generated using two different methods. Method one involved using the crystal structure of the bacterial SemiSWEET transporter to build a homology model. Method two involved using AlphaFold to generate the heterodimeric model of MPC.

The structures of bacterial SemiSWEET transporters have been solved previously via X-ray studies and adopt different conformational states including an inward-facing (IF), occluded (O), and outward-facing (OF) conformations ^[31]^. Because the structure of MPC has not yet been experimentally validated, it is unknown if MPC takes on different conformational states as well. It is suspected MPC’s structure is similar to the bacterial SemiSWEET protein for several reasons. Firstly, it is common for inner-mitochondrial membrane carriers, generally of the SLC25A gene family of solute carriers, to have six transmembrane regions, and the bacterial SemiSWEET homodimer also has six transmembrane domains ^[54], [55]^. Secondly, if MPC1 and MPC2 both had an equal number of transmembrane domains, their N- and C-termini would be on the same side of the membrane. This is important, because it is likely a requirement for the established MPC BRET assays to work efficiently.

Focusing on the OF conformation, which would occur when the transporter initially binds pyruvate or inhibitors in the mitochondrial intermembrane space, the homology model was built from the same conformation of bacterial SemiSWEET transporter (PBD: 4X5N). **Figure 2** shows the resulting structural model of the MPC heterodimer, as well as the structural comparison between the OF and O model hypotheses, to suggest the conformational states that may occur in the hMPC. The positioning of MPC in the membrane is based on the positioning of the bacterial SemiSWEET transporter as well ^[31]^. The predicted transmembrane regions and positioning in the membrane are shown in **Figures 2B & 2C**, as predicted by Desmond and TmAlphaFold.

**Figure 2:**
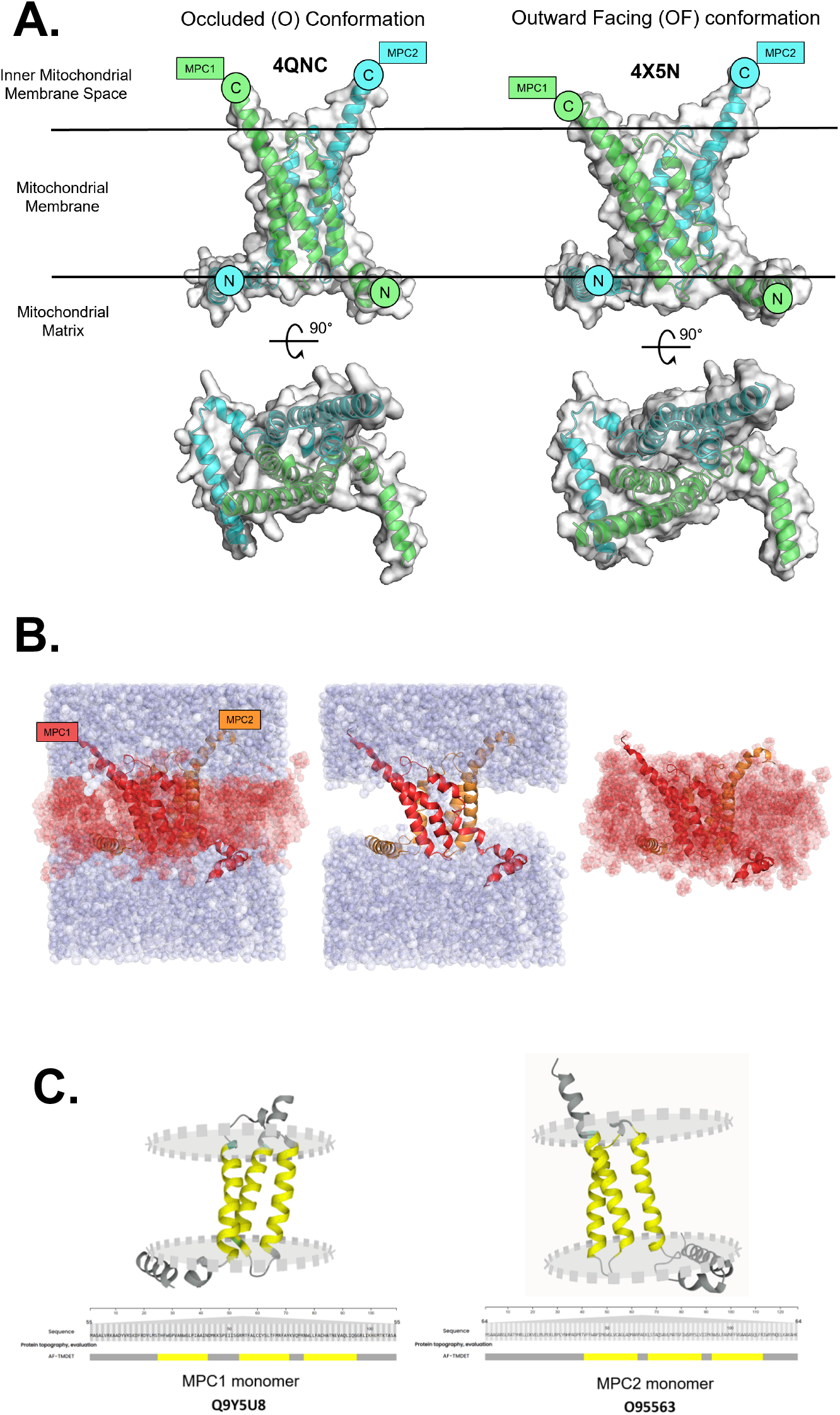
MPC homology model structural prediction matches bacterial SemiSWEET conformations. **A**. Predicted MPC heterodimer structure based on homology modeling with crystal structures of the bacterial SemiSWEET transporter. Two models were generated, one from an occluded (O) (PDB: 4QNC) and the other from an outward-facing (OF) (PDB: 4X5N) conformation of the SemiSWEET protein. Orientation within the mitochondrial membrane was predicted based on SemiSWEET orientation. MPC1 monomer is in green and MPC2 monomer is in blue with the N and C-termini labeled. **B**. Prediction of MPC OF homology model positioning within the membrane predicted by Desmond molecular simulations. Blue represents the aqueous environment and red represents the hydrophobic lipid membrane environment. MPC monomers are colored red (MPC1) and orange (MPC2), respectively. **C**. TmAlphaFold prediction of the transmembrane regions of each MPC monomer, MPC1 (Uniprot: Q9Y5U8) and MPC2 (Uniprot: O95563). Yellow represents regions predicted to fall within the membrane. The grey discs represent the boundaries of the membrane, and the grey represents regions of the protein that do not fall within the membrane.

The other method used AlphaFold to generate the heterodimeric structural models. **Figure 3** details the models generated using this method, after Protein Prep by Schrödinger. AlphaFold is unable to distinguish between potential different conformations, unlike the homology models which are built upon that hypothesis. However, when comparing the generated models with the bacterial SemiSWEET transporters, and calculating binding pockets and tunnels, these models sample a range of conformational states, including OF, IF, or O conformations. Topologies are consistent with the placement of the N-terminal tails on the mitochondrial matrix side of the inner mitochondrial membrane ^[3]^. Based on our prediction of membrane placement shown in **Figure 2B**, it appears that these N-terminal helices reside along the membrane interface, so it is possible their placement is mostly dependent on the lipid membrane, which is not present in our models, and has less impact on the overall transport function. This is potentially supported by a previous study in that a hypomorphic mutant involving N-terminal truncation of the first 16 amino acids of murine MPC2 (~3/4 of the N-terminal helix) results in only ~25% reduction in pyruvate oxidation rates and very mild phenotype that was likely driven more so from reduced expression levels than reduced pyruvate transport function ^[9]^. Therefore, for these models, the overall structure of the transporter’s core was focused on, as opposed to the N-terminal helix positioning.

**Figure 3:**
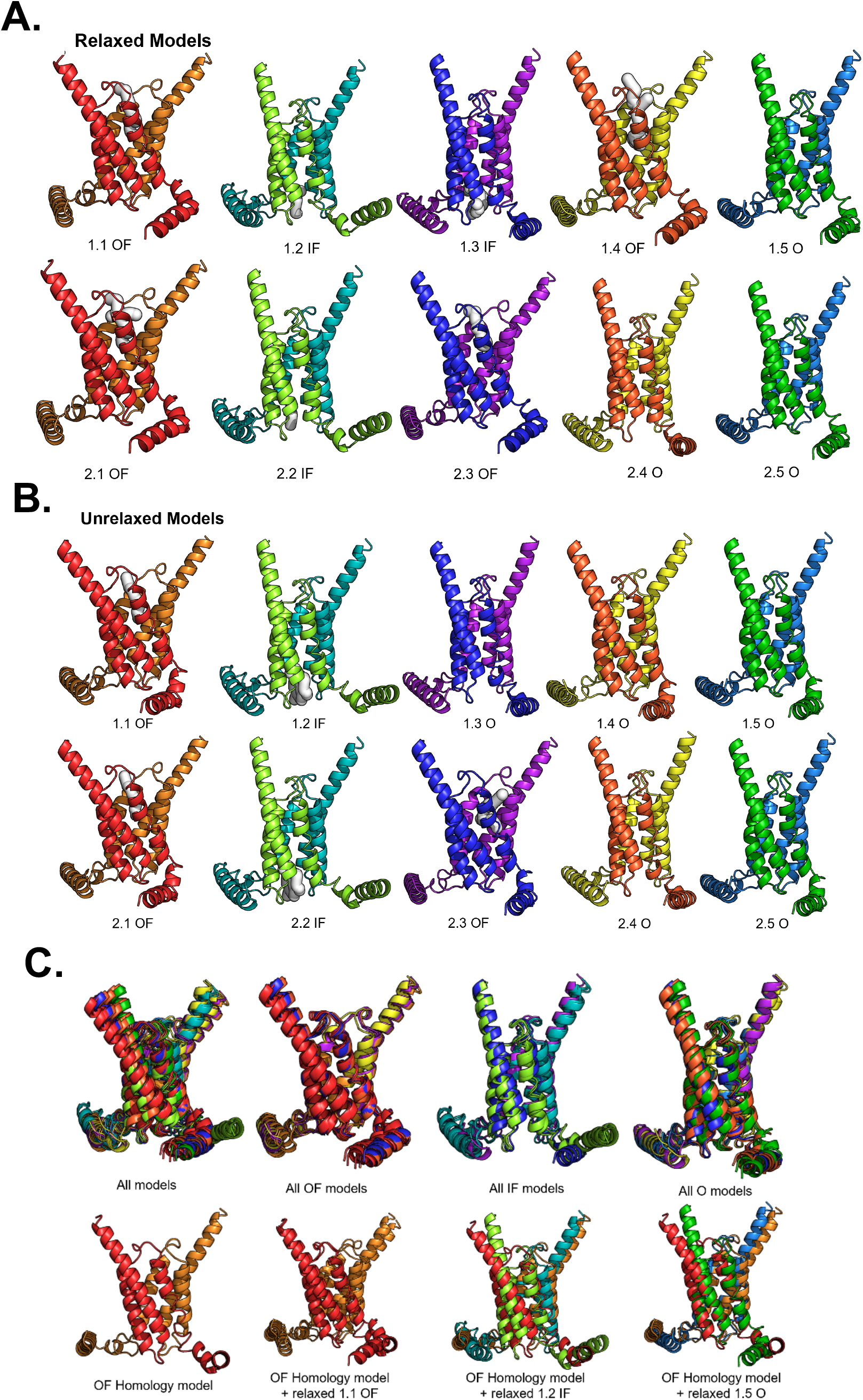
Schrödinger-modified Alpha-Fold models of the MPC heterodimer complex sample different conformational states. **A**. MPC heterodimer complex relaxed models generated using AlphaFold2 multimer v3 software, modified by Schrödinger Protein Prep, and relaxed with Amber. Internal pockets predicted by *MOLEonline* are shown in white. **B**. The similar models generated using AlphaFold2 multimer v3 and Schrödinger Protein Prep, in unrelaxed conditions. Internal pockets predicted by *MOLEonline* are shown in white. Models are categorized into the OF, IF, or O conformation based on *MOLEonline*. **C**. Superposition of all models, plus overlap of the models in each conformation. Representation on how closely the AlphaFold predicted models match the homology model prediction, grouped by the three conformations as well. The two MPC monomers are represented with different colors in each model with the MPC1 monomer on the left and the MPC2 monomer on the right.

MOLE*online* ^[42]^ was used to identify the tunnels and cavities in the models. These comparisons of AlphaFold-generated models are shown in **Figure 3**, along with the pockets identified by MOLE*online*. Conformational identity was determined based on if the model had an internal upper core (OF), lower core (IF), or no core (O) as revealed by MOLE*online*. Of note, comparing the conformations when the models were relaxed versus unrelaxed largely did not alter the conformational identities. The two exceptions were that model 1.3 when relaxed was IF, but changed to O when unrelaxed, while model 1.4 when relaxed was OF, but changed to O when unrelaxed **(Figure 3A-B)**. The IF and O conformation identities were further supported by discoveries during the generation of ligand-binding pockets, discussed in more detail below.

### Docking of known MPC inhibitors

The potential binding pockets in the MPC heterodimer were generated by SiteMap **(Figure 4)**. All the AlphaFold generated models predicted regions fell within five different sites shown in **Figure 4B**. The validity of the predicted regions was determined based on the complex’s predicted positioning in the membrane, as seen in **Figure 2B**. Binding sites 2, 3, and 4 are situated on the outside of the complex at least partially within the membrane region. Because of this placement, it is unlikely those sites would be orthosteric ligand binding sites. However, small parts of the binding pocket may be solvent-exposed and act as an allosteric binding region, so all binding sites were retained for further analysis.

**Figure 4:**
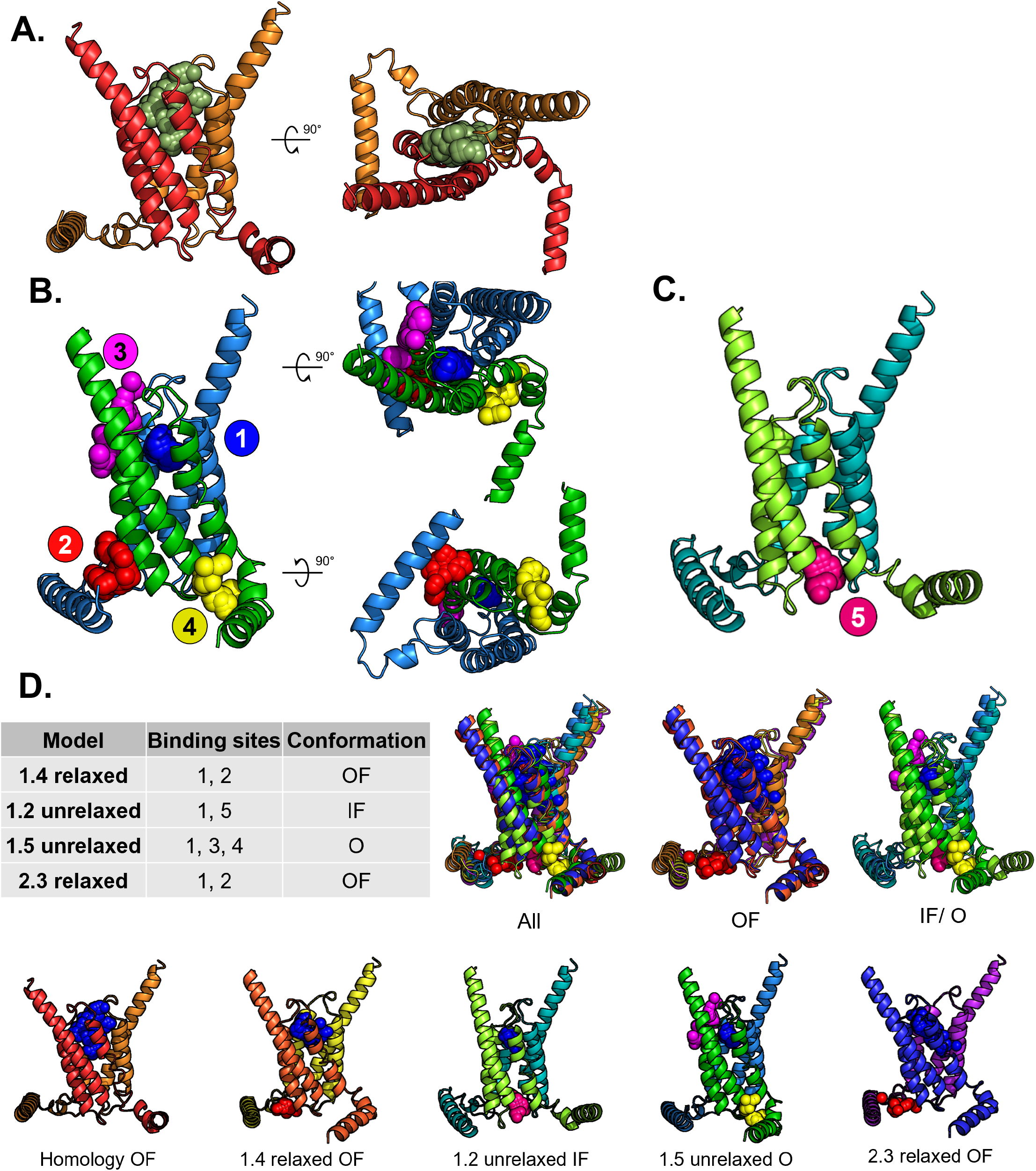
SiteMap binding site predictions in MPC structural models reveal five potential ligand binding regions. **A**. Homology structural model of MPC complex with SiteMap prediction of binding region. This region aligns with site 1 as seen in Figure 4B. **B**. In the Schrödinger-prepped AlphaFold predicted models, SiteMap predicted binding regions in five different areas. The first four sites are modeled on structural model O 1.5 unrelaxed and are color coded (Site 1 = blue, site 2 = red, site 3 = magenta, site 4 = yellow). **C**. A fifth binding site region was identified by SiteMap in two of the AlphaFold predicted models, IF unrelaxed 1.2 and 2.2, shown here in pink. This region is in the center middle of the core as opposed on the outside edges like sites 2 and 4. **D**. Four of the 21 structural models were chosen for future studies based on their broad coverage of different conformations (OF, IF, and O), the five different binding sites, and different AlphaFold generation parameters. The chosen models are the 1.4 relaxed OF, 1.2 unrelaxed IF, 1.5 unrelaxed O, and 2.3 relaxed OF. Their structures and binding site regions are shown here, along with overlap between all the models, just the models in the OF conformation, or just the models in the IF/O conformation.

**Figure 4C** shows a rarer fifth binding pocket region that appeared in two of the IF conformation models, seeming to be the bottom matrix-side of the open transporter channel only accessible in this conformation. Therefore, it is possible that these two models may be the only ones truly in an IF conformation, whereas those lacking binding site 5 are in an O conformation.

Four of the AlphaFold models were chosen for further analysis because they best encompass the different OF and IF conformations of the MPC heterodimer, as well as the five potential binding sites for docking. The chosen structural models were the OF homology model, OF 1.4 relaxed, IF 1.2 unrelaxed, O 1.5 unrelaxed, and OF 2.3 relaxed, as shown in **Figure 4D**. The numbering of these models is based on the parameters used to generate the AlphaFold structures. Group 1 did not use a reference database during generation, but group 2 used the PDB as a reference database. Relaxed models had their energy states relaxed by the program, whereas unrelaxed did not. Receptor grids were generated for each of the structural models and the different binding pockets. While there certainly may be limitations or inaccuracies from docking to unrelaxed models, these two models were included as binding sites 4 and 5 were not identified in any of the relaxed models.

Once the binding pockets were defined, Glide was used to dock known MPC ligands and inhibitors and compare their generated extra precision (XP) Glide scores. Pyruvate (the endogenous ligand), UK-5099 (one of the strongest known experimental MPC inhibitors and gold standard), and several TZDs (known MPC inhibitors and PPARγ agonists), listed in **Table 1**, were investigated. There are several potential approaches to inhibiting the MPC and pyruvate transport. One could be an inhibitor binding allosterically and locking MPC in either the IF or OF conformation so it cannot undergo the gated conformational changes allowing transport. Another approach could be the inhibitor entering and competitively blocking the pyruvate binding site. A third approach could be finding an inhibitor that binds to the interior side of MPC and lies over the core opening, blocking pyruvate transport into the matrix. This blockage would likely be on the inner mitochondrial membrane space side as opposed to the matrix side, based on the compound having to initially enter this space. All possibilities were kept in mind as inhibitors were docked. The results of this docking to the various MPC structural models in the different binding pocket regions are reported in **Table 2** with averaged Glide scores.

**Table 2:** Average Glide docking score of each ligand to each MPC structural model binding site.

| Ligand | MPC Model | Conformation | Binding site | Glide score average |
| --- | --- | --- | --- | --- |
| Pyruvate | 1.4 relaxed | OF | 1 | -3.714 |
|  |  |  | 2 | -2.992 |
|  | 1.2 unrelaxed | IF | 1 | -2.837 |
|  |  |  | 5 | -4.931 |
|  | 1.5 unrelaxed | O | 1 | -2.821 |
|  |  |  | 3 | -2.770 |
|  |  |  | 4 | -3.733 |
|  | 2.3 relaxed | OF | 1 | -3.205 |
|  |  |  | 2 | -3.216 |
| UK-5099 | 1.4 relaxed | OF | 1 | -6.096 |
|  |  |  | 2 | -5.218 |
|  | 1.2 unrelaxed | IF | 1 | N/A |
|  |  |  | 5 | -5.946 |
|  | 1.5 unrelaxed | O | 1 | N/A |
|  |  |  | 3 | -4.310 |
|  |  |  | 4 | -5.514 |
|  | 2.3 relaxed | OF | 1 | -7.840 |
|  |  |  | 2 | -3.178 |
| Rosiglitazone | 1.4 relaxed | OF | 1 | -4.755 |
|  |  |  | 2 | -3.098 |
|  | 1.2 unrelaxed | IF | 1 | N/A |
|  |  |  | 5 | -5.162 |
|  | 1.5 unrelaxed | O | 1 | N/A |
|  |  |  | 3 | -3.184 |
|  |  |  | 4 | -3.745 |
|  | 2.3 relaxed | OF | 1 | -5.860 |
|  |  |  | 2 | -3.307 |
| Ciglitazone | 1.4 relaxed | OF | 1 | -4.099 |
|  |  |  | 2 | -3.254 |
|  | 1.2 unrelaxed | IF | 1 | N/A |
|  |  |  | 5 | -5.764 |
|  | 1.5 unrelaxed | O | 1 | N/A |
|  |  |  | 3 | -3.058 |
|  |  |  | 4 | -3.291 |
|  | 2.3 relaxed | OF | 1 | -5.840 |
|  |  |  | 2 | -3.183 |
| Pioglitazone | 1.4 relaxed | OF | 1 | -5.438 |
|  |  |  | 2 | -2.654 |
|  | 1.2 unrelaxed | IF | 1 | -0.900 |
|  |  |  | 5 | -6.089 |
|  | 1.5 unrelaxed | O | 1 | N/A |
|  |  |  | 3 | -3.817 |
|  |  |  | 4 | -4.371 |
|  | 2.3 relaxed | OF | 1 | -6.960 |
|  |  |  | 2 | -3.576 |
| **Troglitazone** | 1.4 relaxed | OF | 1 | -5.993 |
|  |  |  | 2 | -3.506 |
|  | 1.2 unrelaxed | IF | 1 | N/A |
|  |  |  | 5 | -6.821 |
|  | 1.5 unrelaxed | O | 1 | N/A |
|  |  |  | 3 | -3.382 |
|  |  |  | 4 | -4.025 |
|  | 2.3 relaxed | OF | 1 | -5.859 |
|  |  |  | 2 | -2.982 |
| **Mitoglitazone** | 1.4 relaxed | OF | 1 | -2.863 |
|  |  |  | 2 | 1.283 |
|  | 1.2 unrelaxed | IF | 1 | N/A |
|  |  |  | 5 | -4.124 |
|  | 1.5 unrelaxed | O | 1 | N/A |
|  |  |  | 3 | -2.607 |
|  |  |  | 4 | -1.149 |
|  | 2.3 relaxed | OF | 1 | -3.676 |
|  |  |  | 2 | -1.208 |
| **7ACC2** | 1.4 relaxed | OF | 1 | -7.268 |
|  |  |  | 2 | -4.964 |
|  | 1.2 unrelaxed | IF | 1 | N/A |
|  |  |  | 5 | -5.18 |
|  | 1.5 unrelaxed | O | 1 | N/A |
|  |  |  | 3 | -4.731 |
|  |  |  | 4 | -4.925 |
|  | 2.3 relaxed | OF | 1 | -6.546 |
|  |  |  | 2 | -3.122 |
| **Azemiglitazone** | 1.4 relaxed | OF | 1 | -5.257 |
|  |  |  | 2 | -2.681 |
|  | 1.2 unrelaxed | IF | 1 | N/A |
|  |  |  | 5 | -5.509 |
|  | 1.5 unrelaxed | O | 1 | N/A |
|  |  |  | 3 | -2.895 |
|  |  |  | 4 | -5.228 |
|  | 2.3 relaxed | OF | 1 | -7.986 |
|  |  |  | 2 | -2.866 |

The inhibitors that bound each structural conformational model, along with their Glide scores, are reported in **Table 3**. Visualization of the binding pockets with docked inhibitors can be seen in **Figure 5**. All the structural models, regardless of conformation, shared a binding pocket region in the main center of the heterodimer, which was labeled as site 1. Other binding regions were limited by the structure of the models. For example, site 5 was only seen in the IF conformation and demonstrates an opening of the heterodimer on the matrix side of the membrane. Based on the 3D models, it appears that binding sites 1 and 5 are most viable because they fall within the central internal pocket of the protein heterodimer and are not obstructed by the membrane, like the other binding sites 2-4. Binding region 2, as seen in **Figure 5B**, shows the ligand Pioglitazone entirely resting on top of the N-terminal helix, nestled in a space very likely occupied by the membrane, meaning that is unlikely to be a biologically relevant binding pocket as it is inaccessible. Although parts of these regions close to the core could be allosteric sites where an inhibitor could bind and then cover the core, preventing pyruvate from transporting through the heterodimer, the investigated ligands usually sat fully within the inaccessible region. It is possible that to fully investigate the allosteric pocket potential, a model including the membrane would need to be generated.

**Table 3:**
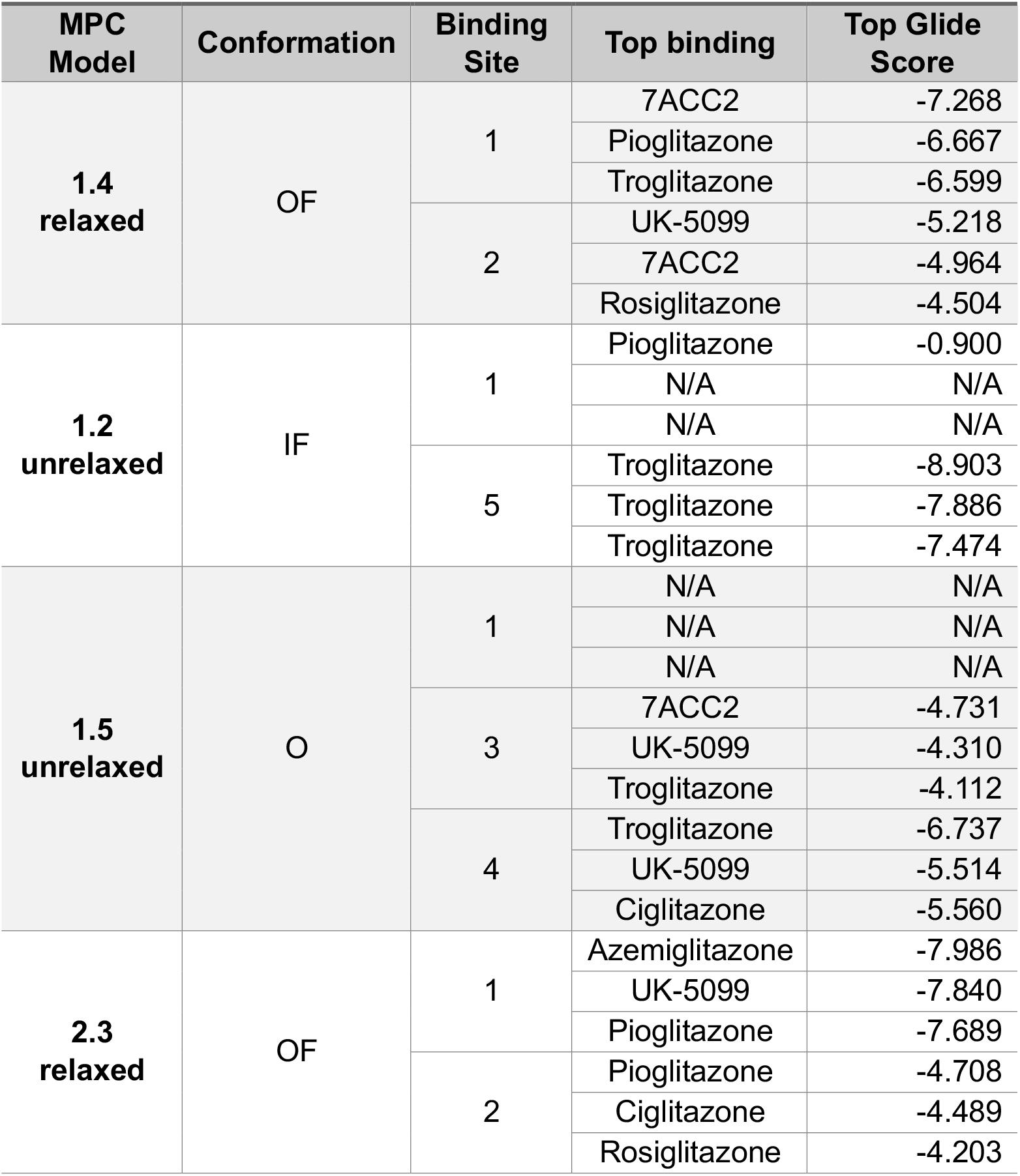
Top docking ligands for each MPC structural model binding site with Glide docking scores.

**Figure 5:**
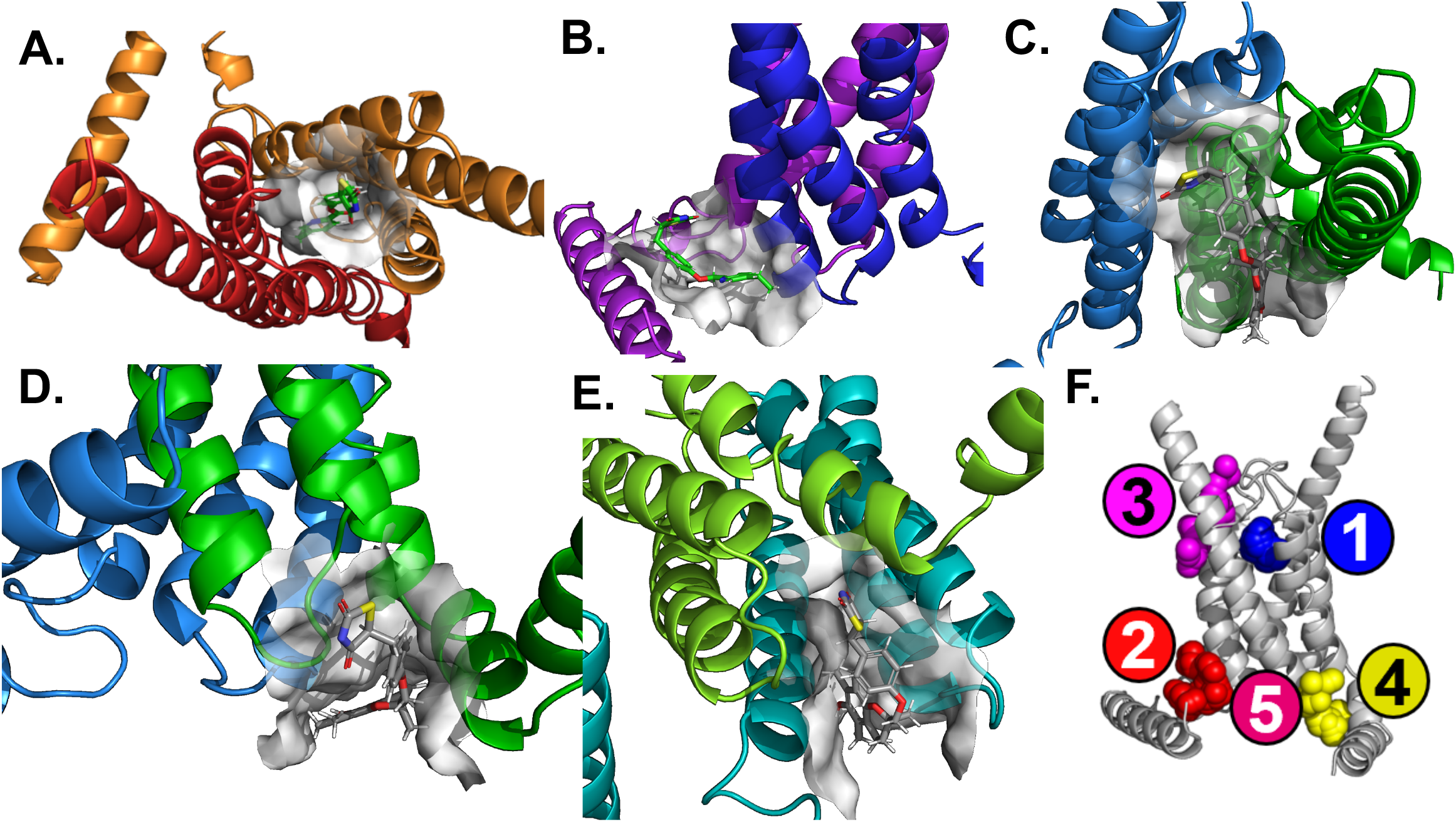
Certain predicted MPC binding sites are likely invalid. **A**. Mitoglitazone docked to binding site 1 in the OF homology model. **B**. Pioglitazone docked to binding site 2 in OF 2.3 relaxed model. **C**. Troglitazone docked to binding site 3 in O 1.5 unrelaxed model. **D**. Troglitazone docked to binding site 4 in O 1.5 unrelaxed model. **E**. Troglitazone docked to binding site 5 in IF 1.2 unrelaxed model. A graphical representative of the different binding sites mapped to the MPC complex allows for orientation of the different three-dimensional models in space. **F**. Representation of physical locations of all 5 binding sites.

Binding regions 1 and 5, depending on the conformation, were able to achieve the best Glide scores, suggesting that those binding regions will be most biologically relevant, especially because they align with the hypothesized MPC model of transport (a central core, the heterodimer flexes between the OF and IF to transport pyruvate through the core). The best Glide scores were achieved by Troglitazone binding to the IF 1.2 unrelaxed conformation in binding region 5 and three different inhibitors (UK-5099, Pioglitazone, and Ciglitazone) binding to the OF 2.3 relaxed in binding region 1. These all achieved a Glide score of −7 or less, with a −10 score indicating optimal free energy for binding, suggesting these Glide scores to be quite good. Two of the models, IF 1.2 unrelaxed binding region 1 and O 1.5 unrelaxed binding region 1, were unable to dock all of the inhibitors. In general, UK-5099, Troglitazone, and Pioglitazone seemed to have the highest Glide scores, which aligns with current experimental data that UK-5099 is a potent MPC inhibitor.

It is important to note that pyruvate had poor Glide scores despite being an endogenous ligand. This is likely for two reasons. First, the transport of pyruvate is transient, so it may not strongly bind to the protein, resulting in poor Glide scores from docking. Second, because pyruvate is such a small molecule compared to the other compounds, it will naturally have fewer sites for interactions, computationally resulting in worse Glide scores. The resulting Glide scores also imply that the bigger TZDs that were unable to bind externally were also too large to get into the core and block it. Therefore, it is questionable if binding site 5 is a viable region to target with inhibitors, because the compounds would likely be too large to go through MPC’s core, unless they entered the mitochondrial matrix via a different pathway first.

Although SiteMap identified binding region 1 for the IF conformation, only pioglitazone with a poor Glide score was able to dock to site 1 in the IF conformation. This is likely because the intermembrane space pocket was closed off in this conformation, preventing the larger inhibitors from accessing and binding. In the context of using these inhibitors as drugs, they would first arrive from the inner mitochondrial membrane space, and if the protein is in the closed-off IF conformation, MPC would be fully inaccessible to them. Therefore, because it is unknown how often and under what conditions MPC flips between the IF and OF conformation, it might be needed to develop an inhibitor that will bind MPC regardless and lock it between conformations.

In general, UK-5099, Troglitazone, and Pioglitazone had the highest Glide scores, making them good potential starting candidates for developing a therapeutic drug. However, the non-specific binding shared with PPARγ first needs to be addressed.

### Chemically important groups for binding to MPC versus PPARγ

Once several compounds with reasonable Glide scores for docking to MPC were identified, the chemical groups of the compounds important for binding to PPARγ were investigated, and those results were compared to the groups needed for binding to MPC. The goal would be to find common chemical motifs that could be modified to remove the PPARγ specificity while maintaining binding to MPC. To do this, the already solved crystal structure of PPARγ in complex with rosiglitazone (PBD: 7AWC) ^[48]^ was used and the same known MPC inhibitors from before, listed in **Table 1**, were docked to the model. The Glide scores from PPARγ are reported in **Table 4**.

**Table 4:** Average Glide docking scores of ligands to PPARγ.

| Ligand | Average Glide Score |
| --- | --- |
| UK-5099 | -9.564 |
| 7ACC2 | -7.493 |
| Rosiglitazone | -12.039 |
| Ciglitazone | -10.796 |
| Pioglitazone | -10.729 |
| Troglitazone | -7.823 |
| Mitoglitazone | -9.901 |
| Azemiglitazone | -12.248 |

Overall, docking to PPARγ resulted in much better Glide scores than docking to the MPC models. TZDs are known for their interaction with PPARγ, so it is unsurprising that their interaction would be stronger with PPARγ over MPC. Of note, these Glide scores do not completely agree with experimentally known potencies for the various TZD molecules on PPARγ binding or transcriptional activation ^[21], [22]^, and these Glide results are likely affected by the PPARγ structure used which was solved in complex with rosiglitazone. Although UK-5099 is functionally different than the TZDs and has not been reported to interact with PPARγ ^[24]^, several studies have indicated that UK-5099 and its analogs can exert similar effects to that of the TZDs ^[56]^. Therefore, it is not entirely surprising that UK-5099 docked to PPARγ with reasonable Glide score. Next, using the ligand interaction task in Schrödinger, interactions within the top docking hits for each inhibitor compound were compared. These interactions with different chemical groups, including hydrogen bonds, salt bridges, pi-pi interactions, and pi-cation interactions between MPC and PPARγ are reported in **Figure 6**. The most common interactions for PPARγ binding involved hydrogen bond formation between the oxygen and nitrogen in the cyclopentane ring while MPC binding involved many other interactions and atoms. Similar interactions showed up during docking to MPC, but many other interactions between different atoms could be focused on, thus providing solid potential for eliminating PPARγ binding to get MPC specificity in a future drug.

**Figure 6:**
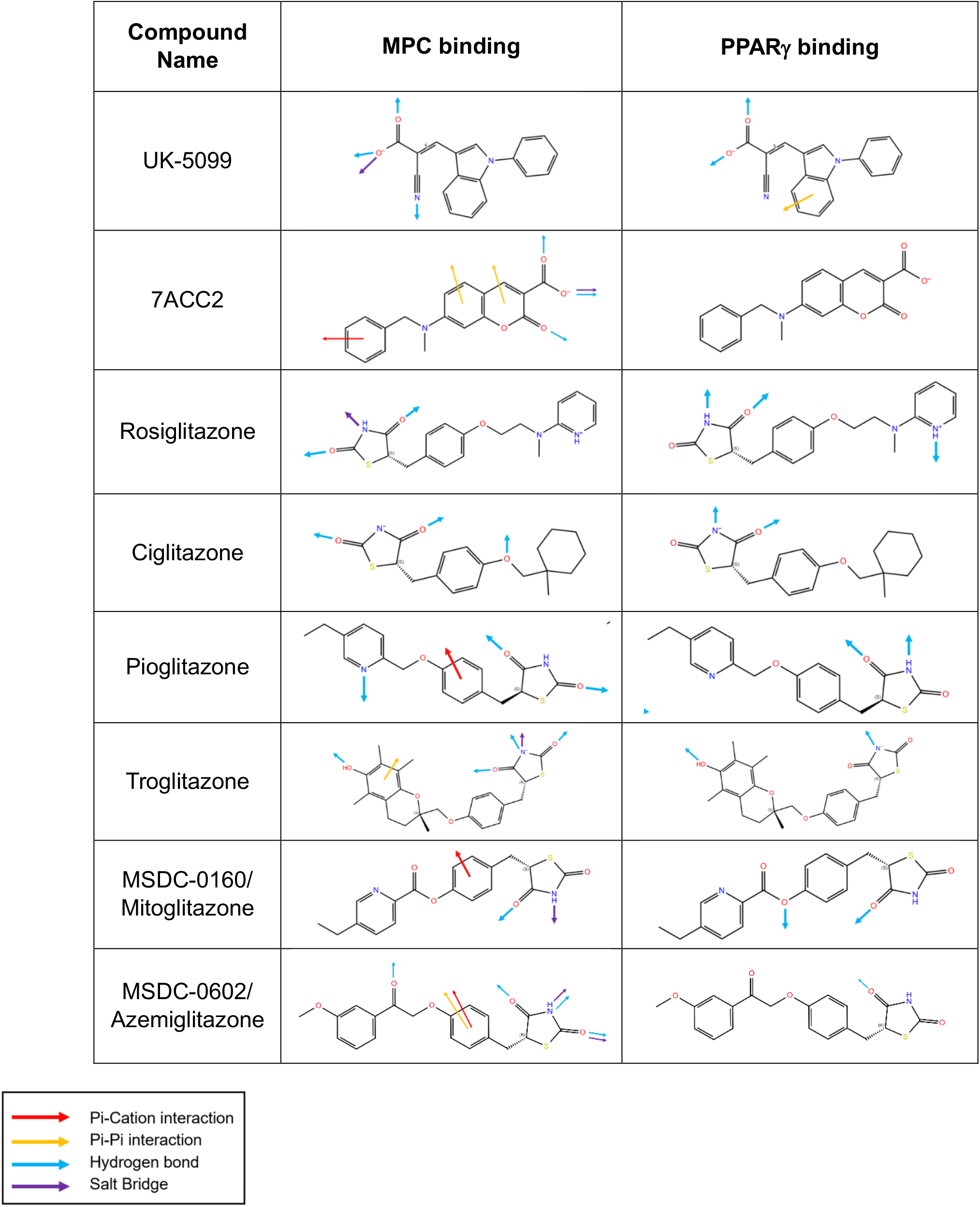
Different chemical group interactions required for MPC binding over PPARγ binding. The following different arrow colors represent different interactions between the docked ligands and proteins. Red = Pi-cation interactions. Yellow = Pi-pi interactions. Blue = hydrogen bond formation. Purple = salt bridge formation. **A**. Interactions for binding between UK-5099, Rosiglitazone, Ciglitazone, Pioglitazone, Troglitazone, Mitoglitazone, Azemiglitazone, and MPC vs. PPARγ.

Previous studies using the aforementioned BRET assay identified three important residues for either binding pyruvate/inhibitors or coupling the binding to conformational changes ^[18], [57]^. These three residues are Phe66 on MPC1 and Asn100 and Lys49 on MPC2. **Figures 7A & 7B** show these three residues highlighted in two of our different structural models: the OF homology model binding site 1 and the IF 1.2 unrelaxed binding site 5. Percentage breakdown of the different MPC amino acids involved in docking interactions in the different IF/O and OF conformational models is shown in **Figure 7C**. Lys49 showed up frequently in docking regardless of conformational model, although it was more prevalently involved in the OF model. While many phenylalanine and alanine residues showed up during docking, Phe66 and Asn100 specifically did not. It is possible that these residues are more important for structure and conformation, allowing inhibitors to bind, or it is possible that the actual core of the MPC is longer, and is impacted by the flexing between conformations, which would not be well reflected in a static model.

**Figure 7:**
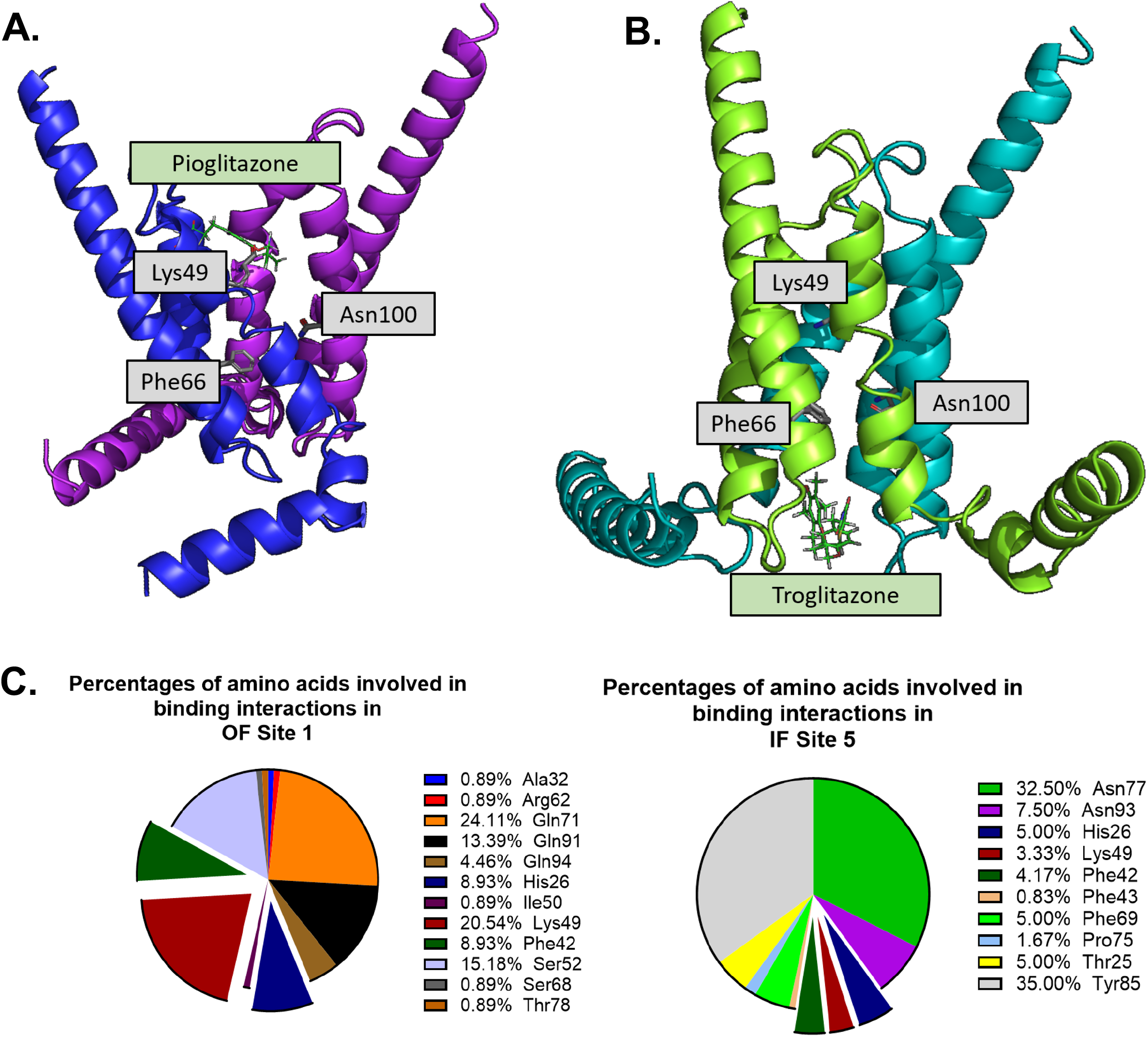
Vital residues imply larger binding pocket than predicted. **A**. Pioglitazone, in green, docked to OF 2.3 relaxed binding site 1. In grey are three previously predicted residues critical for the binding and function of substrates and inhibitors. **B**. Troglitazone, in green, docked to IF 1.2 unrelaxed binding site 5. In grey are three previously predicted residues critical for the binding and function of substrates and inhibitors. These grey residues are Phe66 on MPC1, and Lys49 and Asn100 on MPC2. **C**. Percent representation of how often different MPC amino acids are involved in binding in the different OF and IF models. Pie chart pieces coming out of the graph represent three amino acid residues that were present in each binding model (both OF site 1 and IF site 5).

A previous study from 2021 performed modeling and molecular dynamics simulations of the MPC heterodimer using different modeling software, TrRosetta ^[58]^. As the Rosetta software only accepts a single amino acid sequence, an alanine/glycine linker was inserted between MPC1 and MPC2 to create the heterodimer model which then required manual shifting of the N-terminal amphipathic helices in the previous study. Several results of this previous study were confirmed in this current study, such as the general positioning of the helices and the existence of different conformational states. Their simulations also showed MPC predominately in an IF conformation. Our models revealed a near even split of conformations between OF, IF, and O, while their previous methods did not distinguish between the IF and O conformations, and without that distinction, the prevalence of IF preference is reflected here as well. Their study reported several key residues (Asn33, Leu36, His84, Asn87, in MPC1, and Leu52, Ala55, Leu75, Thr78, Gly79, Trp82, Asn100, and Val103 in MPC2) which UK-5099 interacted with that were not identified by our docking **(Figure 7C)**. These differences are likely due to the advancements made in molecular modeling since this previous study in 2021. The overall agreements between the homology models and the AlphaFold heterodimer predictions from our current study lead us to have increased faith in the accuracy of these models.

## Conclusions

Overall, the models generated in this current study uncovered important foundational knowledge about the potential conformational states of the human MPC, potential binding sites and interactions of different known inhibitors with MPC, and how the interactions chemically differ from PPARγ binding. These results are important for future studies investigating the MPC structure and *in vitro* structural dynamics of the MPC complex, as well as efforts to develop MPC-specific therapeutics for metabolic diseases.

## Acknowledgments

This work was supported by seed grants from Saint Louis University Institute for Drug and Biotherapeutic Innovation (IDBI) and President’s Research Fund (PRF) to K.S.M. C.M.H. is supported by Pharmacological Sciences training grant T32 GM008306, and a Saint Louis University Department of Biochemistry & Molecular Biology Doisy Scholar fellowship.

## Data Availability Statement

The data supporting the findings of this study are available upon reasonable request to the corresponding author.

## Athor Contributions

C.M.H.: study conception, generated models, performed computational analyses, wrote manuscript draft, and edited the manuscript, funding; J.K.W.: assist with computational analyses, and edited the manuscript; R.D.: assist with model generation, study advising, wrote and edited the manuscript; C.A.: study conception, assist with computational analyses, study advising, edited the manuscript; K.S.M.: study conception, funding, study advising, and writing and editing of the manuscript. All authors approved the final version of the manuscript.

